## Supplementary Table 1 for "A Notch-dependent transcriptional mechanism controls expression of temporal patterning factors in *Drosophila* medulla"

**Supplementary Table 1. List of primers used for experiments described.**

| Experiment | Forward Primer | Reverse Primer |
| --- | --- | --- |
| Enhancer bashing GMR35H02 |  |  |
| Cloning R2.2 | TTAGGCGCGCCAGTGCGTGTCTGCCCTTTCATTTTG | ATATATGCGGCCGCCCCAGCTAGCTCCCTTCACTCTTCT |
| Cloning L1.5 | TTAGGCGCGCCGAATCGAAATGCTTCCCCGCCTCG | ATATATGCGGCCGCTGAACGTGCAACATCAAAGGCCGC |
| Cloning M1.0 | TTAGGCGCGCCGCGGCCTTTGATGTTGCACGTTCA | ATATATGCGGCCGCGGACAGTTCGGAATGTGCCTCGA |
| Cloning slpf1 | TTAGGCGCGCCGAATCGAGTGGTGAGCGATAG | ATATATGCGGCCGCTCTGATATTTTTCACGGCTCA |
| Cloning slpf2 | TTAGGCGCGCCTTCGACCTTGTAGTGGCAAG | ATATATGCGGCCGCCGGAGATCGGAAGGTTAGTG |
| Cloning slpf3 | TTAGGCGCGCCTCTCCTTGTTGCTCCTCACA | ATATATGCGGCCGCTGAACGTGCAACATCAAAGG |
| Enhancer bashing d5778 enhancer |  |  |
| d5778 cloning | TTAGGCGCGCCTGGTCTTTTACGTTAATCTGGGCAGCT | ATATATGCGGCCGCACATTACGCATTGCATTCCTCCTCCTT |
| d5778f08 | TTAGGCGCGCCCATTAACTCGAGTCTGGTTTCCGAT | ATATATGCGGCCGCCGTACATATTCTCCAGGAGTTCGGTC |
| Su(H) cloning in pUAST-attB-Dam | ATATATGCGGCCGCAAATGAAGAGCTACAGCCAATTTAATTTAAACGCCGCC | AAAATACTCGAGTCAGGATAAGCCGCTACCATGACTATTCCATTGC |
| CRISPR deletion of u8772 enhancer |  |  |
| u8772 gRNA pCFD5 |  |  |
| Fragment 1 | GCGGCCCGGGTTCGATTCCCGGCCGATGCATCGAAATTTCCTGGTATTCGGTTTTAGAGCTAGAAATAGCAAG | CGCCTCGCCCAAATGCATTTTGCACCAGCCGGGAATCGAACCC |
| Fragment 2 | AAATGCATTTGGGCGAGGCGGTTTTAGAGCTAGAAATAGCAAG | TTAGTTTGATTGTTTCGAAGTGCACCAGCCGGGAATCGAACCC |
| Fragment 3 | CTTCGAAACAATCAAACTAAGTTTTAGAGCTAGAAATAGCAAG | ATTTTAACTTGCTATTTCTAGCTCTAAAAC AAACTGGAAGTCATTGACCCTGCACCAGCCGGGAATCGAACCC |
| Verifying u8772 deletion | TTGCAAATACTTTTTATTCAAGGAATCGAC | AATCTCAAGTTTGGTGTTTGTAATTTTTGG |
| Experiment | **Forward Primer** | **Reverse Primer** |
| CRISPR deletion of d5778 enhancer |  |  |
| d5778 gRNA pCFD5 |  |  |
| Fragment 1 | GCGGCCCGGGTTCGATTCCCGGCCGATGCAATAAGTCCTTGGGTAATACGGTTTTAGAGCTAGAAATAGCAAG | AACATTTATCTAGGACATCTTGCACCAGCCGGGAATCGAACCC |
| Fragment 2 | AGATGTCCTAGATAAATGTTGTTTTAGAGCTAGAAATAGCAAG | TGTTGGCAAGCGGCGCTTCATGCACCAGCCGGGAATCGAACCC |
| Fragment 3 | TGAAGCGCCGCTTGCCAACAGTTTTAGAGCTAGAAATAGCAAG | ATTTTAACTTGCTATTTCTAGCTCTAAAACTTTCGATATCCCAGCTCCTTTGCACCAGCCGGGAATCGAACCC |
| Verifying d5778 deletion | CTATTGAAGGGCGGACATATTAGACAACAATTGGATCGCTTG | CTGCATTCCATCCCGTCGCATCCTTGTC |
| GFP deletion from pJR12 |  |  |
| Fragment 1 | TTAGAGATGCATCTCAAAAAAATGGTGGGCATAATAGTGTTGTTTATATATATCAAAAATAACAAC | CCACCGGTCGCCACCGACGTCAGC GGCCGGCCGC |
| Fragment 2 | GTCGCGGCCGGCCGCTGACGTCGGTGGCGACCGGTGGATCGTTTAAACAGGCC | CAATAACTCGAGGAGCGCCGGAGT ATAAATAGAGGCGCTTCGTCTACG |
| Verifying GFP deletion | CCATTATAAGCTGCAATAAACAAGTTAACAAC | GTCGCTAAGCGAAAGCTAAGC |
| dpn enhancer cloning | TTAGGCGCGCCCTTCGCTTTTGCCTG GTCGGCTCATCGG | ATATATGCGGCCGCACGCCTCGTCCTGGCACCCTC |
| NICD cloning | TATTTAACCGGTTATTATCAAATGTAGATGGCCTCGGAACCCTTG | ATAATAGTTTAAACATGAGTACGCAAAGAAAGCGGGCAC |
| PCR splicing ‘Stop’ cassette with NICD |  |  |
| Fragment 1 | TATTTAACCGGTTATTATCAAATGTAGATGGCCTCGGAACCCTTG | GGAAGTTCCTATTCTCTAGAAAGTATAGGAACTTCGAATTCCAAAATGAGTACGCAAAGAAAGCGGGCAC |
| Fragment 2 | GTGCCCGCTTTCTTTGCGTACTCATTTTGGAATTCGAAGTTCCTATACTTTCTAGAGAATAGGAACTTCC | ATAATAGTTTAAACGAAGTTCCTATTCTCTAGAAAGTATAGGAACTTCCCCGC |
