## Supplementary Table 3 for "A Notch-dependent transcriptional mechanism controls expression of temporal patterning factors in *Drosophila* medulla"

**Supplementary Table 3.** **List of gene blocks used for reporter assays and for cloning Ey coding sequence into the pUAST-Dam-attB plasmid**. Restriction enzyme sites (underlined) and a few overhang bases are added to facilitate cloning.

220-Ey-1d (199 bp)

TTAGGCGCGCCGAATCGAGTGGTGAGCGATAGTTTTCACATGTTTTGTTCGAGCATCGCAACAGATTTGAGCTTTGCACGTAATCAAGTGGAAATGAATTTGACCAACACAAGAGGTTTATACACACATCACATTTTCTGCCTTTATGTTTTTGCGGTGTCCCATTAGTTTGATTGTTTCGAAGGCCACTGAGCCGTGAAAAATATCAGAGCGGCCGCATATT

220-Ey-2m (220 bp)

TTAGGCGCGCCGAATCGAGTGGTGAGCGATAGTTTTCACATGTTTTGTTCGAGCATTAATTCCGTTTAAATGATTTCCGCAACAGATTTGAGCTTTGCACGTAATCAAGTGGAAATGAATTTGACCAACACAAGAGGTTTATACACACATCACATTTTCTGCCTTTATGTTTTTGCGGTGTCCCATTAGTTTGATTGTTTCGAAGGCCACATAGCCGTGAAAAATATCAGAGCGGCCGCATATAT

220-SuH-1-4m (220 bp)

TTAGGCGCGCCGAATCGAGTGGTGAGCGACTAGGAGACGATGTTTTGTTCGAGCATTAATTCCGTTTAAATGATTTCCGCAACAGATTTGAGCTTAGACTGTAATCAAGCGTAAGTGGATTTGACCAACACAAGAGGTTTATACACACATCACATTTTCTGCCTTTATGTTTTTGCGGTGTCCCATTAGTTTGATTGTTTCGAAGGCCACTGAGCCGTAAAGCGAGTCAGAGCGGCCGCATATAT

220-SuH-1m4m (220 bp)

TTAGGCGCGCCGAATCGAGTGGTGAGCGACTAGGAGACGATGTTTTGTTCGAGCATTAATTCCGTTTAAATGATTTCCGCAACAGATTTGAGCTTTGCACGTAATCAAGTGGAAATGAATTTGACCAACACAAGAGGTTTATACACAATCACATTTTCTGCCTTTATGTTTTTGCGGTGTCCCATTAGTTTGATTGTTTCGAAGGCCACTGAGCCGTAAAGCGAGTCAGAGCGGCCGCATATAT

220-Scro-1m (220 bp)

TTAGGCGCGCCGAATCGAGTGGTGAGCGATAGTTTTCACATGTTTTGTTCGAGCATTAATTCCGTTTAAATGATTTCCGCAACAGATTTGAGCTTTGCACGTAATCGTGTGGAAATGAATTTGACCAACACAAGAGGTTTATACACACATCACATTTTCTGCCTTTATGTTTTTGCGGTGTCCCATTAGTTTGATTGTTTCGAAGGCCACTGAGCCGTGAAAAATATCAGAGCGGCCGCATATAT

220-Slp1-1m2m (220 bp)

TTAGGCGCGCCGAATCGAGTGGTGAGCGATAGTTTTCACATGTACCAAGCGAGCATTAATTCCGTTTAAATGATTTCCGCAACAGATTTGAGCTTTGCACGTAATCAAGTGGAAATGAATTGATCGACTCCAAGAGGTTTATACACACATCACATTTTCTGCCTTTATGTTTTTGCGGTGTCCCATTAGTTTGATTGTTTCGAAGGCCACTGAGCCGTGAAAAATATCAGAGCGGCCGCATATAT

Ey gene block for cloning into pUAST-attB-Dam plasmid

AGATCTGCGGCCGCAAATGTTTACATTGCAACCAACTCCAACTGCTATAGGCACCGTGGTTCCCC

CATGGTCAGCGGGAACATTGATAGAGCGCCTGCCGTCTTTAGAAGACATGGCTCACAAGGGTCACAGTGGAGTAAATCAGCTGGGTGGCGTTTTTGTTGGAGGAAGGCCTTTGCCAGATTCAACACGGCAAAAAATTGTCGAACTGGCACATTCTGGAGCTCGGCCATGTGATATTTCTCGAATTCTGCAAGTATCAAATGGATGTGTGAGCAAAATTCTCGGGAGGTATTATGAAACAGGAAGCATACGACCACGTGCTATCGGAGGATCCAAGCCACGTGTGGCCACAGCCGAAGTCGTTAGCAAAATTTCGCAGTACAAACGCGAGTGTCCTAGCATATTTGCTTGGGAAATTCGGGATAGATTACTTCAGGAGAACGTTTGTACTAACGATAATATACCAAGTGTGTCCTCAATAAACCGTGTATTGAGAAACTTGGCTGCGCAAAAGGAGCAGCAAAGCACGGGATCCGGGAGCTCCAGCACATCCGCCGGCAACTCAATCAGCGCAAAAGTGTCTGTCAGCATCGGTGGCAACGTGAGCAATGTGGCAAGCGGATCGAGAGGCACGTTGAGCTCTTCCACCGATCTTATGCAGACAGCCACTCCTCTTAACTCTTCGGAAAGCGGTGGCGCAAGCAACTCCGGGGAGGGTAGTGAACAGGAGGCGATTTACGAGAAGCTTCGGCTGTTAAATACTCAGCACGCTGCAGGACCAGGACCACTGGAGCCTGCCAGAGCAGCGCCCTTGGTAGGTCAATCACCCAACCACCTAGGAACCCGATCCAGCCACCCCCAGCTGGTGCACGGTAACCATCAGGCACTACAGCAGCATCAACAGCAGAGCTGGCCGCCCCGTCACTATTCCGGATCTTGGTACCCCACCTCTCTTAGCGAAATACCCATCTCATCGGCTCCCAATATCGCATCCGTTACGGCGTATGCATCAGGACCTTCACTTGCTCACTCACTGAGTCCACCCAACGACATCGAAAGCCTGGCCAGTATCGGTCACCAGAGAAACTGCCCCGTTGCAACGGAGGACATACATTTAAAAAAAGAACTTGATGGTCATCAGTCCGATGAAACGGGCTCCGGTGAAGGTGAAAACTCCAATGGTGGCGCTTCAAATATAGGAAACACTGAGGATGATCAAGCTCGGCTCATACTAAAAAGAAAGTTGCAACGCAATCGAACATCTTTCACGAACGACCAGATAGACAGTCTTGAAAAAGAGTTTGAACGAACACACTATCCAGATGTTTTTGCCCGCGAACGTTTGGCTGGAAAGATTGGGTTGCCAGAGGCAAGAATTCAGGTTTGGTTCTCAAACCGTCGAGCAAAATGGCGTCGCGAGGAGAAGCTGCGAAACCAGCGAAGAACACCAAATTCCACAGGAGCTAGTGCAACTTCTTCCTCTACATCGGCAACCGCCTCTTTGACTGACAGCCCTAACAGCCTAAGTGCTTGTTCCTCGCTGCTGTCCGGATCAGCTGGGGGTCCCTCAGTCAGTACCATTAATGGCTTATCGTCTCCAAGCACATTGTCTACTAATGTCAATGCTCCAACGCTTGGCGCTGGGATCGATAGCTCTGAAAGCCCAACACCAATCCCGCACATTCGGCCTAGCTGCACCTCTGACAATGACAATGGTCGTCAAAGTGAAGATTGCAGAAGAGTTTGTTCTCCATGCCCACTTGGCGTTGGCGGGCATCAAAATACTCATCATATCCAGAGCAATGGTCACGCCCAAGGTCATGCACTTGTTCCTGCCATTTCGCCACGACTCAATTTTAATAGTGGTAGTTTCGGCGCGATGTACTCCAACATGCATCATACGGCGTTATCCATGAGCGATTCATATGGGGCGGTTACGCCGATTCCGAGCTTTAACCACTCAGCTGTCGGTCCGCTGGCTCCGCCATCGCCAATACCGCAACAGGGCGATCTTACCCCTTCCTCGTTATATCCGTGCCACATGACCCTACGACCCCCTCCGATGGCTCCCGCTCACCATCACATCGTGCCGGGTGACGGTGGCAGACCTGCGGGCGTTGGCCTAGGCAGTGGCCAATCTGCGAATTTGGGAGCAAGCTGCAGCGGATCGGGATACGAAGTGCTATCTGCCTACGCGTTGCCACCGCCCCCTATGGCGTCGAGCTCTGCTGCTGATTCAAGCTTCTCAGCCGCGTCCAGTGCCAGCGCTAATGTGACCCCACATCACACCATAGCCCAAGAATCATGCCCCTCTCCGTGTTCAAGCGCGAGCCACTTTGGAGTTGCTCACAGTTCTGGGTTTTCGTCCGACCCGATTTCACCGGCTGTATCTTCGTATGCACATATGAGCTACAATTACGCGTCGTCCGCTAACACCATGACGCCTTCCTCCGCCAGCGGCACATCAGCACACGTGGCCCCGGGAAAACAACAGTTCTTCGCCTCCTGTTTCTACTCACCGTGGGTCTAGCTCGAGGGTACC

850-Ey-1m2d3d (808 bp)

TTAGGCGCGCCCATTAACTCGAGTCTGGTTTCCGATTCCGATTTCGCTTCCTCCTGCCAACTTATTTCTATATCTTCTCCCTATGTGGCTTGTGTGTGTGAAACAAAAACGTTTGTTTCAATACGTTGGCTTCGTGCATTTTACGGTGTTGGGAAACAGACGAAATGGACTCATAATTGATGTTAAGTGTCTGCCACAGTCGCAGCCGCAAATTCAGTGGCACAACTCCGTCGCAGCCAAATGCCATTTGCTTTTCACATCCAGGTCGAACGGCGTTGCCTTGTTGACTTTGTTTTTGCTACTCATTGCCGCGATTTGGGTTAGGCATGGGGTATGTGCGCACTGTGGGAACTTTGGATTACTCAGATGAAACAGCATTTAGGACACTATGCAGCTGGAAAGATAAACTAGTTGATAGCTACTCATTTACTCATTTACTACTTACTACTAATTTAATGCATTTTTAACAACTTTAAGCTACACAAGCCAAAACTAATGGGTATTTTATAGTCCTATTTAACCCCTTTAACGAATGCATCCTTTTACCTTTTTTGGTCACGGCAGCTGAACTCTGCCCTTTCGTTGGGGGTGACTCCTCCCTCCCGTACTCCCTCCCTCCCTCCCCTCCCTCTCCGCGCCACAGTCGACCTTGTCAAGTACCTTGTTAGCTGTTGGGCAAATGCAGCGGGATCGAAAATAAAAAGCGAAACGCATCGAGAACTTCCCAAGAAAACGGCGAGTCAAAGTTGAGAAAACGCTGCTTCCGTTTAATTGACAATTGAACCCGAACCCGGACCGAACTCCTGGAGAATATGTACGGCGGCCGCATATAT

850-SuH-1m2m3m6m (850 bp)

TTAGGCGCGCCCATTAACTCGAGTCTGGTTTCCGATTCCGATTTCGCTTCCTCCTGCCAACTTATTTCTATATCTTCTCCCTTTGTGCCCTGTGTGTGTGCGACAGATACGTTTGTTTCAATACGTTGGCTTCGTGCATTTTACGGTGTACGGATACAGACGAAATGGACTCATTGATTCCAATTGACTGATTTCAATTGATGTTAAGTGTCTGCCACAGTCGCAGCCGCAAATTCAGTGGCACAACTCCGTCGCAGCCAAATGCCATTTGCTTTTCACATCCAGGTCGAACGGCGTTGCCTTGTTGACTTTGTTTTTGCTACTCATTGCCGCGATTTGGGTTAGGCATGGGGTATGTGCGCACTGTCGGATCTTTGGATTACTCAGATGAAACAGCATTTAGGACACTATGCAGCTGGAAAGATAAACTAGTTGATAGCTACTCATTTACTCATTTACTACTTACTACTAATTTAATGCATTTTTAACAACTTTAAGCTACACAAGCCAAAACTAATGGGTATTTTATAGTCCTATTTAACCCCTTTAACGAATGCATCCTTTTACCTTTTTTGGTCACGGCAGCTGAACTCTGCCCTTTCGTTGGGGGTGACTCCTCCCTCCCGTACTCCCTCCCTCCCTCCCCTCCCTCTCCGCGCCACAGTCGACCTTGTCAAGTACCTTGTTAGCTGTTGGGCAAATGTGCCACACAAGTGGCTCACATCAGCGGGATCGAAAATAAAAAGCGAAACGCATCGAGAACTTCCCAAGAAAACGGCGAGTCAAAGTTCAGATAACGCTGCTTCCGTTTAATTGACAATTGAACCCGAACCCGGACCGAACTCCTGGAGAATATGTACGGCGGCCGCATATAT

850-SuH-1-6m (850 bp)

TTAGGCGCGCCCATTAACTCGAGTCTGGTTTCCGATTCCGATTTCGCTTCCTCCTGCCAACTTATTTCTATATCTTCTCCCTTTGTGCCCTGTGTGTGTGCGACAGATACGTTTGTTTCAATACGTTGGCTTCGTGCATTTTACGGTGTACGGATACAGACGAAATGGACTCATTGATTCCAATTGACTGATTTCAATTGATGTTAAGTGTCTGCCACAGTCGCAGCCGCAAATTCAGTGGCACAACTCCGTCGCAGCCAAATGCCATTTGCTTTTCACATCCAGGTCGAACGGCGTTGCCTTGTTGACTTTGTTTTTGCTACTCATTGCCGCGATTTGGGTTAGGCATGGGGTATGTGCGCACTGTCGGATCTTTGGATTACTCAGATGAAACAGCATTTAGGACACTATGCAGCTGGAAAGATAAACTAGTTGATAGCTACTCATTTACTCATTTACTACTTACTACTAATTTAATGCATTTCTAAGTACTTTAAGCTACACAAGCCAAAACTAATGGGTATTTTATAGTCCTATTTAACCCCTTTAACGAATGCATCCTTTTACCTTTTTTGGTCACGGCAGCTGAACTCTGCCCTTTCGTTGGGGGTGACTCCTCCCTCCCGTACTCCCTCCCTCCCTCCCCTCCCTCTCCGCGCCACAGTCGACCATGTCAAGTACCTTGTTAGCTGTTGGGCAAATGTGCCACACAAGTGGCTCACATCAGCGGGATCGAAAATAAAAAGCGAAACGCATCGAGAACTTCCCAAGAAAACGGCGAGTCAAAGTTCAGATAACGCTGCTTCCGTTTAATTGACAATTGAACCCGAACCCGGACCGAACTCCTGGAGAATATGTACGGCGGCCGCATATAT

850-Scro-1-7d (724 bp)

TTAGGCGCGCCCATTAACTCGAGTCTGGTTTCCGATTCCGATTTCGCTTCCTCCTGCCAACTTATTTCTATATCTTCTCCCTTTGTGCCCTGTGTGTGTGAAACAAAAACGTTTGTTTCAATACGTTGGCTTCGTGCATTTTACGGTGTTGGGAAACAGACGAAATGGACTCATTGATTCCAATTGACTGATTTCAAAGTCGCAGCCGCAAATTCAGTGGCACAACTCCGTCGCAGCCAAATGCCATTTGCTTTTCACATCCAGGTCGAACGGCGTTGCCTTGTTGACTTTGTTTTTGCTACTCATTGCCGCGATTTGGGTTAGGCATGGGGTATGTGCGCACTGTGGGAACTTTGGATTACTCAGATGACAGCTGGAAAGATAAACTAGTTGATAGCTACTCATTTACTCATTTACTACTTACTACTAATTTAATGCATTTTTAACAACTTTAAGCTACATTTATAGTATGCATCCTTTTACCTTTTTTGGTCACGGCAGCTGAACTCTGCCCTTTCGTTGGGGGTGACTCCTCCCTCCCGTACTCCCTCCCTCCCTCCCCTCCCTCTCCGCGCCACAGTCGACCGTTGGGCAAATGTGCCACACAAGTGGCTCACATCAGCGGGATCGAAAATAAAAAGCGAAACGCATCGAGAACTTCCCAAGAAAACGGCGAGTCAAAGTTGAGAAAACGCTGCTTCCGAACCCGGACCGAACTCCTGGAGAATATGTACGGCGGCCGCATATAT

850-Slp1-1d (829 bp)

TTAGGCGCGCCCATTAACTCGAGTCTGGTTTCCGATTCCGATTTCGCTTCCTCCTGCCAACTTATTTCTATATCTTCTCCCTTTGTGCCCTGTGTGTGTGAAACAAAAACGTTTGTTTCAATACGTTGGCTTCGTGCATTTTACGGTGTTGGGAAACAGACGAAATGGACTCATTGATTCCAATTGACTGATTTCAATTGATGTTAAGTGTCTGCCACAGTCGCAGCCGCAAATTCAGTGGCACAACTCCGTCGCAGCCAAATGCCATTTGCTTTTCACATCCAGGTCGAACGGCGTTGCCTTGTTTGCCGCGATTTGGGTTAGGCATGGGGTATGTGCGCACTGTGGGAACTTTGGATTACTCAGATGAAACAGCATTTAGGACACTATGCAGCTGGAAAGATAAACTAGTTGATAGCTACTCATTTACTCATTTACTACTTACTACTAATTTAATGCATTTTTAACAACTTTAAGCTACACAAGCCAAAACTAATGGGTATTTTATAGTCCTATTTAACCCCTTTAACGAATGCATCCTTTTACCTTTTTTGGTCACGGCAGCTGAACTCTGCCCTTTCGTTGGGGGTGActcctccctcccgtactccctccctccctcccctccctctccGCGCCACAGTCGACCTTGTCAAGTACCTTGTTAGCTGTTGGGCAAATGTGCCACACAAGTGGCTCACATCAGCGGGATCGAAAATAAAAAGCGAAACGCATCGAGAACTTCCCAAGAAAACGGCGAGTCAAAGTTGAGAAAACGCTGCTTCCGTTTAATTGACAATTGAACCCGAACCCGGACCGAACTCCTGGAGAATATGTACGGCGGCCGCATATAT
