## Supplementary File 1 for "A Notch-dependent transcriptional mechanism controls expression of temporal patterning factors in *Drosophila* medulla"

Sequence of fragment deleted by CRISPR-Cas9 targeting the u8772 220bp enhancer (5’ -> 3’)

2L: 3817278-3817675

cgcctcgcccaaatgcatttgaatcgagtggtgagcgatagttttcacatgttttgttcgagcattaattccgtttaaatgatttccgcaacagatttgagctttgcacgtaatcaagtggaaatgaatttgaccaacacaagaggtttatacacacatcacattttctgcctttatgtttttgcggtgtcccattagtttgattgtttcgaaggccactgagccgtgaaaaatatcagagaaataaaaatcgaaataccgaaatgagctggttttttgttagcgaaagtgcagatttttcaggactcgcaaagggatgtgattgaagatcttcaggatatttcagcacatgcatcgatatttgtcccaaactggaagtcattgacccagttactttt

Sequence of fragment deleted by CRISPR-Cas9 targeting the d5778 850bp enhancer

2L: 3842548-3844867

aagtccttgggtaatacgaggagaacaaaaacaagtaaggataatattttcataggcaaaacaattgtggttgtgcaaggaagtgacttttgggaacgggaggcacttgcagctgcccaagtatatccaccatatcccatatcctttttccgagtccctaatttcctgggaaagctttcggtgcagttcgccagccaaaacacttgagcacttaaaaaggcgcattaactcgagtctggtttccgattccgatttcgcttcctcctgccaacttatttctatatcttctccctttgtgccctgtgtgtgtgaaacaaaaacgtttgtttcaatacgttggcttcgtgcattttacggtgttgggaaacagacgaaatggactcattgattccaattgactgatttcaattgatgttaagtgtctgccacagtcgcagccgcaaattcagtggcacaactccgtcgcagccaaatgccatttgcttttcacatccaggtcgaacggcgttgccttgttgactttgtttttgctactcattgccgcgatttgggttaggcatggggtatgtgcgcactgtgggaactttggattactcagatgaaacagcatttaggacactatgcagctggaaagataaactagttgatagctactcatttactcatttactacttactactaatttaatgcatttttaacaactttaagctacacaagccaaaactaatgggtattttatagtcctatttaacccctttaacgaatgcatccttttaccttttttggtcacggcagctgaactctgccctttcgttgggggtgactcctccctcccgtactccctccctccctcccctccctctccgcgccacagtcgaccttgtcaagtaccttgttagctgttgggcaaatgtgccacacaagtggctcacatcagcgggatcgaaaataaaaagcgaaacgcatcgagaacttcccaagaaaacggcgagtcaaagttgagaaaacgctgcttccgtttaattgacaattgaacccgaacccggaccgaactcctggagaatatgtacgctgctatccggcatagtcgagtcatcccaagtcatgcgcttaatttcccgtttaaaacgccattcattcaattaagcgaatgaatttgtggcacggcagacgacagcagaagtttttttttcctgaaagaagactatcatcattgagatgccccgagatctctcggctggagctccagatccgatgcgatccaatcggatgagatgagatcgtatggaataggatggagcatgcgggtctctggggtctgtggtctttcgatctttttgtctgcccccgggggctttcttcagtaatcagacgcgggtcaaaatatttaccacttgaccctattgatttaattaaatagtttcaagacattcaactgtttgttggccgcgcaaaatgaatttcgtgtgcgagataactcagatacagtagctgcagtagctgatcaactatctttcagatatgcgcacatttctctctcgttttgtctccgagctgtcaacacagattccaaactgcagacgtgttaattaacgacagagttaactaattgttgttagcaaaatattttcgtaattcgatactaaattcgagttccgctcaacttgcttgcttgtgggcttttgtgttgaacagcacaagacctcaaggaacagtggaacagtgcttcaaagtggggaaaaacatcttataatctagatcatttttaaatttcataaagtctttgattgaaataaaacgtttaagtgaggcaagttcgtattattattagagaaagataactaacctttttggtttatatttaggtcttgaagcactgttcctgctgttttcgctgttgactgcccatgaaacgcttaactgcgagtggcaattggatacctggctgaatccaaatacgaatcttaatctgaatctgcgagtctttgtggccaatgaacacggcagcggcacaacacaaaaacttagcagatactcatgttttatttgtgcatttcgcgcgcgcgttccacttggaaatgctctgtggcaccaaaaggagccactgatgtccaaaatcaaaatggtttcagttcgccggggcaatgcctaaaatgtttctttgtttttgtgttccctgttggcaagcggcgcttcaacagatacagatagatgtgaaatttgttgctaaaaaaaaaagtaagtccataaatcaaatgcctttaaatattcatgagtcaggacaatgtgtgaacaagcgaaaggagctgggatatcgaa
